# Perturbative formulation of general continuous-time Markov model of sequence evolution via insertions/deletions, Part IV: Incorporation of substitutions and other mutations

**DOI:** 10.1101/023622

**Authors:** Kiyoshi Ezawa, Dan Graur, Giddy Landan

## Abstract

**Background:** Insertions and deletions (indels) account for more nucleotide differences between two related DNA sequences than substitutions do, and thus it is imperative to develop a stochastic evolutionary model that enables us to reliably calculate the probability of the sequence evolution through indel processes. In a separate paper (Ezawa, Graur and Landan 2015a), we established the theoretical basis of our *ab initio* perturbative formulation of a continuous-time Markov model of the evolution of an *entire* sequence via insertions and deletions along time axis. In other separate papers (Ezawa, Graur and Landan 2015b,c), we also developed various analytical and computational methods to concretely calculate alignment probabilities via our formulation. In terms of frequencies, however, substitutions are usually more common than indels. Moreover, many experiments suggest that other mutations, such as genomic rearrangements and recombination, also play some important roles in sequence evolution.

**Results:** Here, we extend our *ab initio* perturbative formulation of a *genuine* evolutionary model so that it can incorporate other mutations. We give a sufficient set of conditions that the probability of evolution via both indels and substitutions is factorable into the product of an overall factor and local contributions. We also show that, under a set of conditions, the probability can be factorized into two sub-probabilities, one via indels alone and the other via substitutions alone. Moreover, we show that our formulation can be extended so that it can also incorporate genomic rearrangements, such as inversions and duplications. We also discuss how to accommodate some other types of mutations within our formulation.

**Conclusions:** Our *ab initio* perturbative formulation thus extended could in principle describe the stochastic evolution of an *entire* sequence along time axis via major types of mutations.

[This paper and three other papers (Ezawa, Graur and Landan 2015a,b,c) describe a series of our efforts to develop, apply, and extend the *ab initio* perturbative formulation of a general continuous-time Markov model of indels.]

## Introduction

The evolution of DNA, RNA, and protein sequences is driven by mutations such as base substitutions, insertions and deletions (indels), recombination, and other genomic rearrangements (*e.g.*, Graur and Li 2000; Gascuel 2005; Lynch 2007). Thus far, analyses on substitutions have predominated in the field of molecular evolutionary study, in particular using the probabilistic (or likelihood) theory of substitutions that is now widely accepted (e.g., Felsenstein 1981, 2004; Yang 2006). However, some recent comparative genomic analyses have revealed that indels account for more base differences between the genomes of closely related species than substitutions (*e.g*., Britten 2002; Britten et al. 2003; Kent *et al*. 2003; The International Chimpanzee Chromosome 22 Consortium 2004; The Chimpanzee Sequencing and Analysis Consortium 2005). It is therefore imperative to develop a stochastic model that enables us to reliably calculate the probability of sequence evolution via mutations including insertions and deletions.

Traditionally, the computation of probabilities of indels has been based on hidden Markov models (HMMs) or transducer theories (see, *e.g*., Rivas 2005; Bradley and Holmes 2007; Miklós et al. 2009). However, these methods have two fundamental problems, one regarding the theoretical grounds and the other regarding the biological realism. (See the “background” section in part I (Ezawa, Graur and Landan 2015a) for more details on these problems.)

As an unprecedented approach to these problems on the probabilistic models of indels, we proposed an *ab initio* perturbative formulation of a *genuine* stochastic evolutionary model, which describes the evolution of an *entire* sequence via indels along the time axis (Ezawa, Graur and Landan 2015a). Such a *genuine* evolutionary model is devoid of the aforementioned problems from the beginning. More specifically, our *genuine* evolutionary model is a general continuous-time Markov model of sequence evolution via indels. It allows any indel rate parameters including length distributions, but it does not impose any unnatural restrictions on indels. In part I of this series of study (Ezawa, Graur and Landan 2015a), we gave the theoretical basis of our *ab initio* formulation. Especially, we derived a sufficient and nearly necessary set of conditions under which the probability of an alignment is factorable, like a sort of HMM. In part II (Ezawa, Graur and Landan 2015b), we developed some analytical techniques for performing concrete perturbation analyses. In part III (Ezawa, Graur and Landan 2015c), we developed an algorithm that can calculate the first-approximate probability of each local MSA delimited by gapless columns, given an input MSA and under a given parameter setting including a phylogenetic tree.

Mainly in order to avoid unnecessary confusions of the readers, these studies (Ezawa, Graur and Landan 2015a,b,c) dealt with indels alone. In terms of frequencies, however, substitutions are usually more common than indels (*e.g*., Lunter 2007; Cartwright 2009). Moreover, many experiments suggest that still other mutations, such as genomic rearrangements and recombination, also play some important roles in the evolution of protein and DNA sequences (*e.g.*, Graur and Li 2000; Lynch 2007; Gu et al. 2008). Thus, a natural question arises as to whether the methods and conclusions obtained in these papers are still valid, at least to some degree, even when we consider other types of mutations as well.

In this study, we extend our *ab initio* parturbative formulation of a *genuine* evolutionary model so that it can incorporate other mutations, such as substitutions and genomic rearrangements. In Section 1 of Results & Discussion, we consider the model of sequence evolution via both indels and substitutions. We give a sufficient set of conditions that the probability of an alignment under this model is factorable into the product of an overall factor and the contributions from local evolutionary processes. We also show that, under a set of conditions, the probability can be factorized into two components. One is the “basic” component given by the indel processes alone, and the other is the residue component that concerns the substitution processes alone. Our set of conditions is more general than the commonly used conditions that the indigenous and inserted residue frequencies are equal to the equilibrium frequencies of the substitution model (*e.g*., Thorne et al. 1991, 1992; Miklós et al. 2004; Rivas and Eddy 2008). In Section 2, we show that our *ab initio* formulation of a genuine evolutionary model can be extended to incorporate genomic rearrangements, especially inversions and duplications. In Section 3, we discuss how we can accommodate some other types of mutations within our formulation. Appendix gives mathematical details on Section 1.

This paper is part IV of a series of our papers that documents our efforts to develop, apply, and extend the *ab initio* perturbative formulation of the general continuous-time Markov model of sequence evolution via indels. Part I (Ezawa, Graur and Landan 2015a) gives the theoretical basis of this entire study. Part II (Ezawa, Graur and Landan 2015b) describes concrete perturbation calculations and examines the applicable ranges of other probabilistic models of indels. Part III (Ezawa, Graur and Landan 2015c) describes our algorithm to calculate the first approximation of the probability of a given MSA and simulation analyses to validate the algorithm. Finally, part IV (this paper) discusses how our formulation can incorporate substitutions and other mutations, such as duplications and inversions.

This paper basically uses the same conventions as used in part I (Ezawa, Graur and Landan 2015a). Briefly, a sequence state *s* (∈ *S*) is represented as an array of sites, each of which is either blank or equipped with some specific attributes. And each indel event is represented as an operator acting on the bra-vector, **〈** *s|*, representing a sequence state. More specifically, the operator 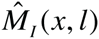 denotes the insertion of *l* sites between the *x* th and (*x* +1) th sites, and the operator 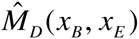 denotes the deletion of a sub-array between (and including) the *x*_*B*_ th and the *x*_*E*_ th sites. See Section 2 of part I for more details. And, as in part I, the following terminology is used. The term “an indel process” means a series of successive indel events with both the order and the specific timings specified, and the term “an indel history” means a series of successive indel events with only the order specified. And, throughout this paper, the union symbol, such as in *A* ∪ *B* and 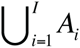, should be regarded as the union of *mutually disjoint* sets (*i.e.,* those satisfying *A* ∩ *B* = ∅ and *A*_*i*_ ∩ *A*_*j*_ = ∅ for *i* ≠ *j* (∈ {1,…, *I*}), respectively, where ∅is an empty set), unless otherwise stated.

## Results & Discussion

### 1. Incorporation of substitutions

For clarity, we focused on the description of indel processes and omitted substitutions in the bulk of the paper. Actually, it is not so difficult to incorporate substitutions into our framework. For this purpose, we first extend the sequence state space *S* (= *S^I^*, *S^II^*, *or S^III^*) so that each site of the sequence will accommodate a residue in the set Ω (see, *e.g*., Subsection 2.1 of part I (Ezawa, Graur and Landan 2015a)). When *S* = *S^I^*(≅ N_0_≡ {0,1, 2,…}), the extended space is 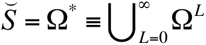, in which each site is assigned a residue *ω* ∈ **Ω**. When *S* = *S^II^*(⊂ ϒ^*^), the ancestry assigned to each site, *υ* ∈ϒ, is replaced with a pair, (*υ*, *ω*) ∈ϒ ×**Ω**. Thus, the extended space is 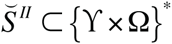. Similarly, when *S* = *S^III^* (⊂ {N_0_ × N_1_}*), the extended space is 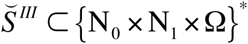, in which each site is assigned a trio, (*σ*, *ξ*, *ω*) ∈ N_0_× N_1_×Ω.(Here *σ* is the source identifier of the inserted (or initial) subsequence, and *ξ* is the relative coordinate within the inserted (or initial) subsequence.) An extended sequence state 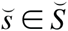 will sometimes be represented as 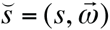 to explicitly show that it is composed of a “basic” component *s* ∈ *S* and a residue component 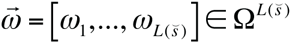. (Here 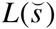 is the length of the sequence 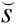.) Once we extended the sequence state space, we can now extend the indel rate operator 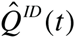 (given in Eqs.(2.4.2a-d) of part I). This is done in two steps. (1) We add the substitution component of the rate operator, 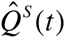. The operator is a generalization of the one given in Eq.(1.2.1’) of part I to a sequence with multiple sites and to a more general substitution model. And (2) we extend the indel component 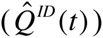 to take explicit account of the residue dependence of indels, including the relative probabilities among residue states filling in each inserted array of sites. See Appendix A1 for details. Then, the total rate operator of the entire evolutionary model 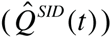 is given by adding the indel and substitution rate operators:

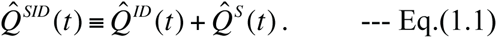

At least formally, we could apply the perturbation expansion technique to the stochastic evolution operator of the entire model, 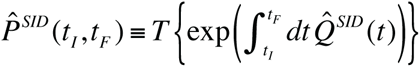 (*T* {…} denotes the time-ordered product.) We can do this by decomposing 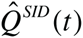 into two parts: 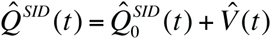, where 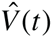 is treated as a “perturbation” part. There are two major ways of doing this. (A) To decompose it into 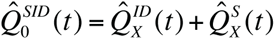 and 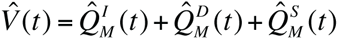. And (B) to decompose it into 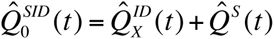 and 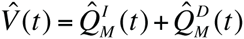. Here 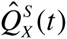 and 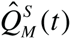 are the exit-rate part and the single-mutation part, respectively, of 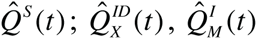; and 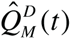, are the exit-rate part, the single-insertion part and the single-deletion part, respectively, of 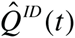.

By mainly using the decomposition (A) and by following the same line of arguments as in Sections 3 and 4 of part I, we can show that the probability of an alignment (whether it is a PWA or a MSA) is factorable into the product of an overall factor and the contributions from regions that can potentially accommodate local evolutionary histories, this time including both indels and substitutions. We showed the factorability under the following sufficient set of conditions. (I) In each region, the rate of each of the substitutions and indels is independent of the portions of the extended sequence states in the other regions. (II) In each region, the increment of the total exit rate, 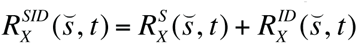, due to each of the substitutions and indels is independent of the portions of the extended states in the other regions. And, if considering a MSA, (III) the factorization of the root sequence state probability (*i.e*., an extended version of Eq.(4.2.8) in part I) holds. See Appendix A2 for details on the derivation.

The factorization of the probability into the local contributions would definitely be helpful. However, it would be at least equally useful to factorize the probability under the entire evolutionary model into the “basic component” and the “residue” component.” Here, the “basic component” is based only on possible indel histories, and the “residue component” is based only on possible substitution histories (and initial residue state distributions). Mainly using the aforementioned decomposition (B), we can show that such an “indel-substitution factorization” can indeed be carried out on the alignment probability if the following four conditions are satisfied. (i) The indel rates are independent of the residue states. (ii) Each finite-time evolution probability of the residue state via substitutions is factorable into the product of site-wise substitution probabilities. (iii) The residue state spectrum of each inserted sub-sequence is factorable into the product of site-wise residue probabilities over the inserted sites. And (iv) the site-wise inserted residue probabilities at each site, {*p*_*I*_ (*ω*; *υ _j_*, *t*)} _*ω*∈Ω_, where *υ _j_* is the ancestry of the site, should satisfy the equation:

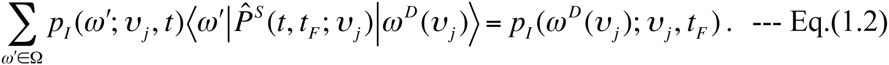

Here 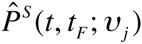 is the site-wise stochastic evolution operator via substitutions. When dealing with a MSA, the following equation also needs to be satisfied:

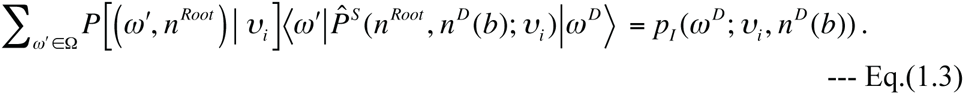

Here, *P*[(*ω* **′**, *n^Root^*)|*υ_i_*] is the probability of the residue *ω* **′** at the site with ancestry *υ _i_* in the sequence at the root node (*n^Root^*). See Appendix A3 for details on the proof of the “indel-substitution factorization” under these conditions. Eq.(1.2) means that the inserted residue frequencies must evolve according to the site-wise stochastic evolution via substitutions. And Eq.(1.3) requires that, at every point along the tree, they should coincide with the residue frequencies that would have evolved through substitutions beginning with {*P*[(*ω*, *n^Root^*)|*υ _i_*]}_*ω* ∈Ω_ at the root. These equations are a generalized version of the following commonly assumed pair of conditions. (a) The residue frequencies are equilibrium frequencies, {*π* (*ω*)}_*ω* ∈Ω_, that satisfy the detailed-balance conditions, 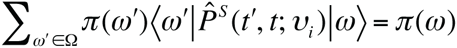. And (b) the inserted residue frequencies must coincide with these equilibrium frequencies (see, *e.g*., Thorne et al. 1991, 1992; Miklós et al. 2004; Rivas and Eddy 2008).

In order to pursue the biological realism further, however, the aforementioned conditions (i)-(iv) would be too restrictive, even though the conditions were somewhat relaxed compared to the common practice of imposing (a) and (b) (instead of (iv)). It may be relatively easy to relax the conditions (ii) and (iii) so that the substitution processes could depend on the residue states of neighboring sites, by, *e.g*., introducing codon models (*e.g*., Yang 2006) (but see also, *e.g*., Lunter and Hein 2004; Arndt and Hwa 2005). The violation of the condition (i) might be tackled at least partially, if the indel rates depend on the residue states only locally, through some motifs that sparsely scatter along the sequence. Specifically, in such a case, we may first factorize the probability into the product of local contributions and then perform the “indel-substitution factorization” on the contributions from regions that are likely devoid of such motifs. The violation of the condition (iv) may be more prevalent and serious, especially for large-scale insertions (see, *e.g*., Waterhouse and Russell 2006; Morgante et al. 2007; Chalopin et al. 2015). Recently, analytical methods where developed for examining the effects of deviation of the inserted residue composition from the substitution-inherent residue composition (Lèbre and Michel 2010, 2013). It may be worth trying to apply some of their methods to the situations where the condition (iv) is violated. Meanwhile, some recent data analyses showed that the substitution rate increases near the sites of indels (Tian et al. 2008; De and Babu 2010). If desired, such effects could be represented in our extended theoretical formulation (described in Appendix A4 in more details), and might be handled similarly to how the violation of the aforementioned condition (i) could be remedied. It remains to be seen whether the remedial methods suggested here actually work, or otherwise whether our formulation could be modified or further extended somehow to efficiently deal with the more biologically realistic features mentioned above.

### 2. Incorporation of inversions and duplications

At least formally, the theoretical formulation developed in this paper can be extended to incorporate other genomic rearrangements (*e.g*., Gascuel 2005; Gu et al. 2008). Here, we discuss the incorporation of inversions (*e.g*., Kelshner and Wendel 1996; Graham et al. 2000) and duplications (*e.g*., Bailey and Eichler 2006; Lynch 2007; Ezawa et al. 2011), as two most important examples.

To incorporate inversion processes, it is convenient to extend the space state, especially *S^II^* or *S^III^*, to accommodate the complement of each site. Specifically, we let a superscript “ *C*” indicate that the state is of a site on the complementary strand of the site before inversion. For example, *υ^C^* denotes the ancestry of the site complementary to a site with the ancestry *υ* (in the space *S^II^*). And (*σ*,*ξ*)*^C^* denotes the attributes of the site complementary to a site with attributes (*σ*,*ξ*) (in the space *S^III^*). And we consider that the complement of the complement is the original: (*X^C^*)*^C^* = *X*. To incorporate duplication processes, we also introduce another indicator, *χ*, telling that the site is on the *χ* th copy of the subsequence. For example, *υ*.*χ* represents the ancestry of the *χ* th copy of the original site with the ancestry *υ* (in the space *S^II^*), and (*σ*,*ξ*, *χ*) represents the attribute of the *χ* th copy of the original site with the attribute (*σ*,*ξ*) (in the space *S^III^*). The state spaces formed by extending the spaces *S^II^* and *S^III^* this way will be represented by *S^IIe^* and *S^IIIe^*, respectively. [NOTE: As for the state space *S^I^*, we do not need to extend it to accommodate inversions and duplications; we just invert or duplicate a sub-array of *unlabeled* sites of a sequence, as well as the *residue states* filling in the sub-array.]

An inversion event could be depicted by an inversion operator, 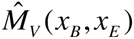, which inverts a sub-array of sites between (and including) the *x*_*B*_ th and *x*_*E*_ th sites.

For example, the action of 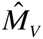 (2,4) on the basic state

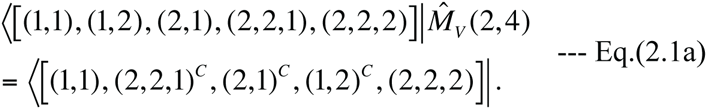

In principle, we could also define the inversion of a region sticking out of either sequence end. For example, the action of 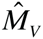 (0, 2) on the above state could be represented by something like the following:

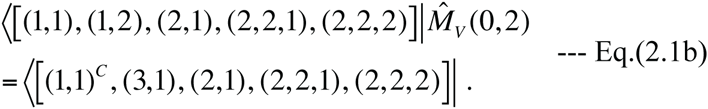

The point is that this operation replaces the 2nd site with the complement of the 0th site. The 0th site was outside of the region under consideration, and we formally assigned it a new attribute, (3,1)*^C^*, before inversion. Thus, an inversion sticking out of either sequence end is effectively equivalent to a simultaneous operation of a deletion and an insertion, and also a smaller-scale inversion when the inverted region within the sequence is longer than the region sticking out. The inversion rate operator, 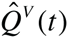, is also defined similarly to how the indel rate operator is defined via Eqs.(2.4.1a,b) of part I (Ezawa, Graur and Landan 2015a):

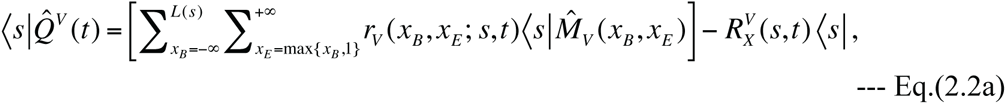

with the exit rate:

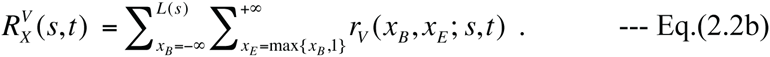

If the inversion rates, *r*_*V*_ (*x*_*B*_, *x*_*E*_; *s*, *t*), are space-homogeneous, the exit rate will be an affine function of the sequence length and the probability of a LHS equivalence class of inversion processes will be factorable. Unfortunately, inversions are known to occur preferentially on palindrome sequences or between inverted repeats (*e.g*., Kelshner and Wendel 1996; Gu et al. 2008). Nevertheless, even if we take account of such structural dependence, the probability may be more or less factorable. This is because a simple inversion does not change the sequence length or much of the inverted repeat structure, and thus because 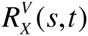 is expected to change little due to inversions.

A duplication event could be depicted via a duplication operator, 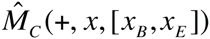 or 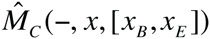, which copies the sub-array between (and ncluding) the *x*_*B*_ th and *x*_*E*_ th sites and inserts the copy between the *x* th and (*x* +1)th sites. The “ +” and “ −” in the 1st argument represent the insertion on the original and complementary strands, respectively. For example, the actions of 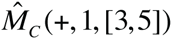 and 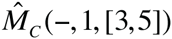 on the above state could be expressed as:

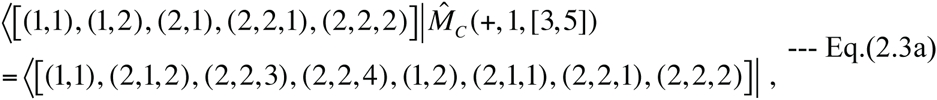

and

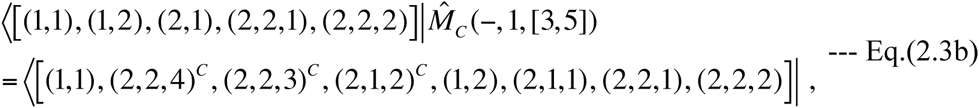

respectively. Again in principle, we could also define the duplication of a region sticking out of either sequence end. For example, the action of 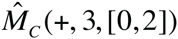 on the above state could be depicted as:

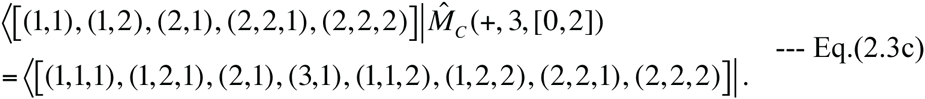

Again, the 0th site was out of consideration before the event, and thus was assigned a new attribute, (3,1), when inserted. In general, the duplication of a region sticking out of either sequence end is effectively equivalent to the simultaneous operation of a duplication of the region within the sequence and an adjacent insertion of a new subsequence. Furthermore, we could define the duplication of a region totally out of the sequence under consideration, *i.e*., *r*_*C*_ (*ε*, *x*, [*x*_*B*_, *x*_*E*_]) with *ε* ∈ {+, −} and with either *x*_*B*_ ≤ *x*_*E*_ **<** 1 or *L*(*s*) **<** *x*_*B*_ ≤ *x*_*E*_. Under the current formulation, its effect is indistinguishable from that of the insertion, *r*_*I*_ (*x*, *x*_*E*_ − *x*_*B*_ +1). Using these duplication operators, the duplication rate operator, 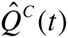, could be defined as:

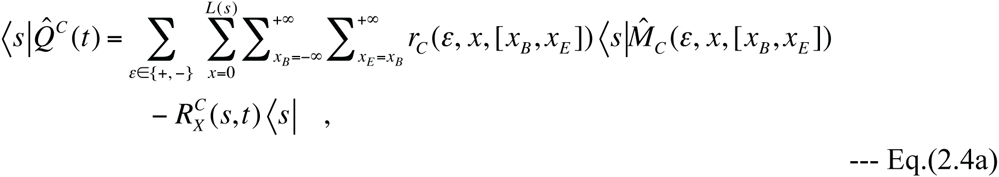

with the duplication exit rate:

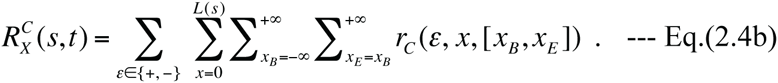

If the duplication rates, *r*_*C*_ (*ε*, *x*, [*x*_*B*_, *x*_*E*_]), are space-homogeneous, 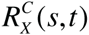 should ecome an affine function of the sequence length, and the probability of a LHS equivalence class of duplication processes could be factorable. But, again, it is unlikely that the duplication rates are space-homogeneous, because duplications preferentially occur between direct repeat motifs (*e.g*., Bailey and Eichler 2006; Gu et al. 2008). It remains to be seen whether the probability is still factorable or not even after taking account of this factor. But some situations with interspersed repeat motifs may be modeled as in Eqs.(5.3.2a,b,c) in part I, at least to some degree, and thus the probabilities may be partially factorable.

Some transposon insertions (*e.g*., Morgante et al. 2007; Chalopin et al. 2015) are essentially duplications (via copy-and-paste mechanisms). Thus, in some cases it might be beneficial to regard the transposons as duplicated using the formulation explained here, rather than handling them as simple insertions. On the other hand, the latter could be done in principle by associating particular sets of residue configurations with elevated insertion rates, via the theoretical formulation briefed in Section 1. Their description as duplications would be beneficial especially when two or more transposons belonging to the same family were inserted into positions that are close to each other, because they could induce secondary genomic rearrangements.

In a traditional alignment (PWA or MSA), an inversion should manifest itself as a pair of equally long gapped regions, interpreted as a deletion and an adjacent insertion, and it will normally be penalized twice. Moreover, commonly used aligners will ignore the fact that the reverse complement of the inverted region can be well aligned with its original region. Meanwhile, a duplication event should cause a gap in a normal alignment, and will be interpreted as an insertion. However, unless we take account of the fact that the duplicated region is in fact homologous to, and thus can be aligned with, its original region, the resulting alignment could often be erroneous. Therefore, by taking explicit account of inversions and deletions when reconstructing an alignment, its accuracy might improve substantially. This kind of attempts has only a short history (see, *e.g*, Paten et al. 2008b, and references therein). The theoretical formulation briefed in this subsection may be conducive to the development of likelihood-based (or Bayesian-based) alignment programs that also take account of genomic rearrangements other than indels. To do so, the rate parameters, such as *r*_*V*_ (*x*_*B*_, *x*_*E*_; *s*, *t*) and *r*_*C*_ (*ε*, *x*, [*x*_*B*_, *x*_*E*_]), will have to be estimated accurately. It would probably be difficult to estimate them directly from the sequence data to be analyzed, because the events are relatively rare. Thus, it would be helpful to estimate the parameter distribution, or relative rates as functions of the length of the duplicated/inverted region, the distance between the original region and the copy-insertion point, residue states of the flanking regions, etc., by analyzing genome-wide data from a large sample of organisms (or individuals from populations). Although the broad dependences of the duplication frequency on the original-copy distance and the orientation were examined in the past (*e.g*., Ezawa et al. 2011), more extensive and thorough analyses will be necessary.

At the population genetic level, genome rearrangements are observed as genomic structural variations (SVs), including copy number variations (CNVs) (*e.g*., Teshima and Innan 2012; Ezawa et al. 2013). In some aspects, population genetics could be regarded as molecular evolution on a very short time scale. Thus, the theoretical formulation unfolded in this paper may be applicable, possibly via some modifications, to the analyses of SVs as well.

### 3. Other mutational mechanisms

Mechanisms of genomic mutations are not limited to substitutions and genome rearrangements. Among the most important ones would be recombination (*e.g*., Saitou and Kitano 2013) and gene conversion (*e.g*., Chen et al. 2007; Ezawa et al. 2010; Fawcett and Innan 2013). Out of them, inter-locus gene conversion (*e.g*., Ezawa et al. 2010; Fawcett and Innan 2013) may be depicted via something similar to the duplication operator proposed in Section 2, but with a modification. Specifically, instead of inserting a sub-sequence, a gene-conversion operator must make a sub-sequence replace another region that is usually paralogous. This description will work as long as the interacting regions are both within a single sequence to be analyzed. However, recombination between alleles (*e.g*., Saitou and Kitano 2013), including inter-allelic gene conversion (*e.g*., Chen et al. 2007), is a mechanism involving two orthologous sequences, and thus the above description will not be naïvely applicable to it. The description will not be applicable to inter-locus gene conversion, either, if one of the interacting regions is out of the subject sequence. A possible way to handle these cases would be to allow the phylogenetic tree to take different topologies and/or branch lengths in different regions of the sequence, as implemented in the molecular evolution simulator, Dawg (Cartwright 2005). This measure should work especially when the alignment probability is factorable.

The copy number change of short tandem repeats, or microsatellites, is another important mutational mechanism (*e.g*., Ellegren 2004; Sainudiin et al. 2004). However, the evolution of microsatellites would be quite refractory to a naïve analysis, because they have quite high mutation rates and show complex mutation patterns (*e.g*., Ellegren 2004; Sainudiin et al. 2004). Thus, unless we are handling very short-term sequence evolution, we should avoid using the perturbation theory developed in this paper. Instead, we should try to solve “exactly,” maybe via a numerical computation, an empirical evolutionary model of microsatellite evolution (*e.g*., Sainudiin et al. 2004). Our current hunch is that, as far as an accurate alignment reconstruction is concerned, a best way would be to remove alleged microsatellites from the sequences before aligning them.

### 4. Boundary conditions and cut-off lengths

In this study, as in the previous studies (Ezawa, Graur and Landan 2015a,b,c), we only considered simple boundary conditions. Each sequence end was either freely mutable or flanked by a biologically essential region that allows no indels. Moreover, the constant cutoff lengths were introduced just for the sake of simplicity, to broadly take account of the effects of various factors that suppress very long rearrangements (such as selection, chromosome size, genome stability, etc.). In real sequence analyses, however, the situations are unlikely to be so simple. (See Discussion of part I (Ezawa, Graur and Landan 2015a) for more details.) In order to pursue further biological realism and to enable further accurate sequence analyses, it would be inevitable to address these issues seriously.

## Conclusions

In a previous study (Ezawa, Graur, and Landan 2015a), we established the theoretical basis of an *ab initio* perturbative formulation of a general continuous-time Markov model, which is a *genuine* stochastic model describing the evolution of an *entire* sequence via indels along the time axis. Then, in two other previous studies (Ezawa, Graur, and Landan 2015b,c), we demonstrated how we can analytically and computationally calculate the probabilities of concrete alignments via our formulation. In these previous studies, we dealt only with insertions/deletions (indels), mainly for clarity of the arguments.

Here in this study, we attempted to incorporate other types of mutations into our *ab initio* perturbative formulation of a *genuine* evolutionary model. We first extended the model to accommodate substitutions. We showed that, under a set of conditions on the model parameters similar to that for the pure indel model, the probability of an alignment is factorable into local contributions also in this extended model. We also gave a set of conditions under which an alignment probability is factorable into the product of an “indel component” and a “substitution component.” We next showed how our evolutionary model can be extended to accommodate two other types of genomic rearrangements, namely, inversions and duplications. We also discussed how to handle other types of mutations such as recombination within the framework of our formulation. Thus, at least in principle, our *ab initio* perturbative formulation could describe the evolution of an *entire* sequence via any major types of mutations along the time axis. It still remains to be seen whether all of these suggested extensions are *computationally* feasible and useful for *practical* sequence analyses, such as the reconstruction of generalized multiple sequence alignments (MSAs). If this proves to be the case, however, our formulation could open up a new venue for the theoretical study of sequence evolution.

## Authors’ contributions

KE conceived of and mathematically formulated the theoretical framework in this paper, implemented the key algorithms, participated in designing the study, performed all the mathematical analyses, and drafted the manuscript. DG and GL participated in designing the study, helped with the interpretation of the data, and helped with the drafting of the manuscript.

## Acknowledgements

This study is dedicated to the late Dr. Keiji Kikkawa, who was a renowned theoretical physicist, one of the key pioneers of the string field theory of the elementary particle physics, and the best ever mentor of K.E. We are grateful to Dr. R. A. Cartwright at Arizona State University for his useful information and discussions that inspired this study. We appreciate the logistic support and the feedback of Dr. Tetsushi Yada at the Kyushu Institute of Technology. We would also like to thank the three anonymous referees of the predecessor manuscript entitled: “Framework that enables approximate lilelihood analysis of insertions/deletions on multiple sequence alignment.” Their comments helped drastically improve the study itself, not to mention the manuscript. This work was a part of the project, “Error Correction in Multiple Sequence Alignments,” which was funded by US National Library of Medicine [grant number LM010009-01 to Dan Graur and Giddy Landan at the University of Houston]. The later stage of this work was also supported by Grants-in-Aid No. 221S0002, which was awarded to Tetsushi Yada by the Ministry of Education, Culture, Sports, Science and Technology of Japan.

## Appendix

Here we give detailed arguments regarding the incorporation of substitutions into our theoretical formulation, as briefly described in Section 1 of Results. We first explain how to extend our formulation to incorporate substitutions in Section A1. Then, in Section A2, we will examine the conditions under which the probability of each alignment is factorable into the product of contributions from regions delimited by preserved ancestral sites (PASs). Next, in Section A3, we will examine the conditions under which the probability of each alignment can be factorized into the “basic” component concerning the indels and the residue component concerning the substitutions (and the initial residue state distribution at each site). Finally, in Section A4, we will discuss how to pursue the biological realism further.

### A1. Formulating sequence evolution via both indels and substitutions

In order to incorporate substitutions into our theoretical formulation of sequence evolution via indels, we first extend the sequence state space *S* (= *S^I^*, *S^II^*, *or S^III^*) so that each site of the sequence will accommodate a residue in the set Ω. (See Subsection 2.1 of part I (Ezawa, Graur and Landan 2015a) for details on the original sate spaces *S^I^*, *S^II^ and S^III^*.) When *S* = *S^I^*(≅N_0_ ≡ {0,1, 2,…}), the extended space is 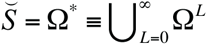, in which each site is assigned a residue *ω* ∈ Ω. When *S* = *S^II^(*⊂ ϒ^*^), the ancestry assigned to each site, *υ* ∈ϒ, is replaced with a pair, (*υ*, *ω*) ∈ϒ ×Ω. Thus, the extended space is 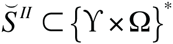. Similarly, when *S* = *S^III^* (⊂ {N_0_× N_1_}*), the extended space is 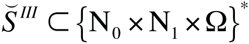, in which each site is assigned a trio, (*σ*, *ξ*, *ω*) ∈ N_0_× N_1_×Ω. (Here *σ* is the source identifier of the initial or inserted (sub)sequence, and *ξ* is the relative coordinate within the initial or inserted (sub)sequence.) An extended sequence state 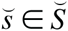 will sometimes be represented as 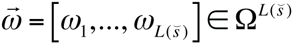 to explicitly show that it is composed of a “basic” component *s* ∈ *S* and a residue component 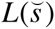. (Here 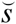 is the length of the sequence state 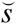.) Once we extended the sequence state space, we can now extend the indel rate operator (defined in Eqs.(2.4.2a-d) of part I). This is done in two steps: (1) adding the substitution component to the rate operator; and (2) extending the indel component to take explicit account of the residue dependence of indels.

First, we add the substitution component of the rate operator, 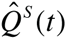, to the indel component, 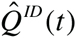 (defined in Eqs.(2.4.2a-d) of part I). This yields the total mutation rate operator:

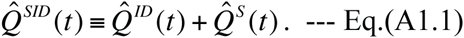

Similarly to the derivation of Eqs.(2.4.2a-d) of part I, the substitution component is written as 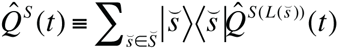 is the substitution rate operator defined on a sequence of length *L*. If we allow only for a single residue substitution at a time, 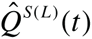 is expanded as: 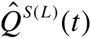, where 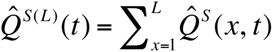 is the operator representing the effect of a substitution at the *x* th site at time *t*. Let 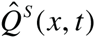 be the operator representing the substitution from the residue *ω* to *ω*′ (≠ *ω*) at the *x* th site. And let 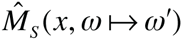 be the residue state at the *x* th site of a sequence with the extended state 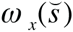. Then, in general, 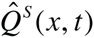 is defined by the following action on the extended sequence states 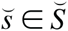 with 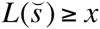:

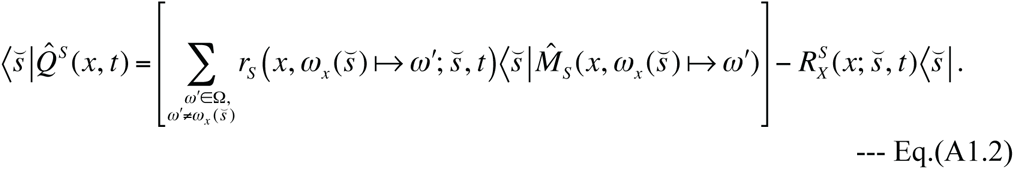

Here the position-specific exit rate, 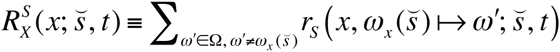 *is* made of the substitution rates at the *x* th site, 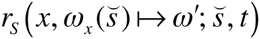, which could in general depend on the extended sequence state, site position, and time. In the special case where all indel rates are zero, the sequence states keep their length and the ancestries *υ_x_* (or attributes (*σ _x_*, *ξ_x_*)) of the sites intact. Thus, the total stochastic evolution operator of the system 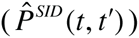 is given solely by its substitution component, 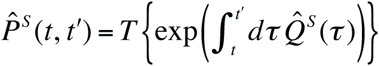. (Here *T* {…} denotes the time-ordered product. See below Eq.(1.1.11) of part I for details.) If desired, we can decompose the substitution rate operator as 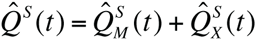. Here, 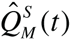 is a linear combination of substitution operators, and 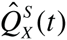 keeps the residue state unchanged while letting the probability decay at the exit rate. Then, in the same line of reasoning as in Subsection 4.1 of part I, the conditional probability 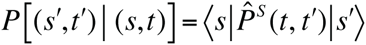 can be factorized into the product of multiplication factors, each of which is contributed from a region that is independent of the others. Actually, in this case, the stochastic evolution operator 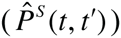 itself becomes factorable into a tensor product, as long as no mutation changes the interdependence structure among the substitution rates at different sites. Especially, when each of the substitution rates depends only on the state at the site of the substitution, *e.g.*, 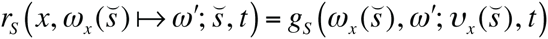, we have:

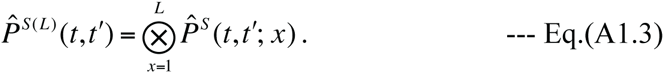

Here, 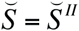, supplemented by Eq.(A1.2), is the stochastic operator describing the evolution of the *x* th site via substitutions during the time interval [*t*, *t***′**]. At least conceptually, Eq.(A1.3) should be familiar to many researchers of molecular evolution (*e.g*., Felsenstein 1981, 2004; Yang 2006).

Second, we extend the indel component of the rate operator, 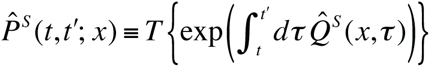 (given in Eqs.(2.4.2a-d) of part I), so that it will take *explicit* account of the possible residue state dependence of indels. First of all, we replace the summations, such as 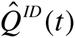 and 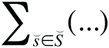, with the extended ones, *i.e*., 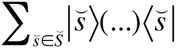 and 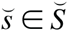. Then, consider the operations of indels on the extended state 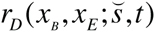, as well as their rates. Regarding deletions, it is easy; we just need to extend the deletion rates *r*_*D*_ (*x*_*B*_, *x*_*E*_; *s*, *t*) to *r*_*D*_ (*x*_*B*_, *x*_*E*_; 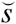, *t*), which explicitly depend also on the residue state component 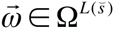 of the extended state 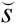. Regarding insertions, however, we need something more. We first introduce a new operator, 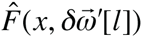, that fills in an array of newly inserted sites, from the (*x* +1) th through the (*x* + *l*) th sites in the post-insertion sequence, with an array of residues, 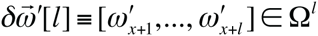. In other words, 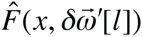 inserts 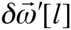 between the *x* th and (*x* +1) th residues of 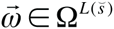. Then, we replace the term *r*_*I*_ (*x*, *l*; *s*, *t*) 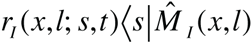 in Eq.(2.4.2b) of part I with 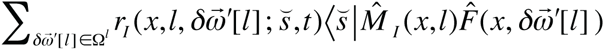. And we also replace *r*_*I*_ (*x*, *l*; *s*, *t*) in the exit rate (Eq.(2.4.1b) or Eq.(2.4.1b’) of part I) with 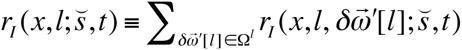. If there is no correlation between the 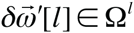 to be inserted and the state 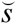 to undergo the insertion, the dependence of the rates on them could be decoupled as 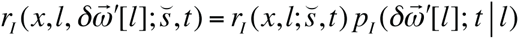. Here 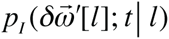 is the probability (or the relative frequency) that 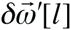 is inserted at time *t*, conditioned on the insertion of *l* sites. (The conditional probabilities satisfy 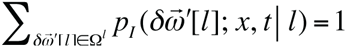. This implies that *r*_*I*_ (*x*, *l*; *s*, *t*) used in the bulk of the paper were something like 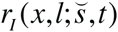 above, considering that the former already *implicitly* depended on the residue states of the sequence before insertion. If the above decoupling holds, and if the dependence on the inserted residues can be factorized as 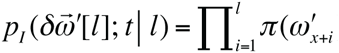, then the inserted residues could be handled as were done in the past (*e.g*., Thorne et al. 1991, 1992; Miklós et al. 2004; Rivas and Eddy 2008). Here *π*(*ω*) is the equilibrium frequency of a residue *ω* (with Σ_ω∈Ω_ *π*(*ω*) = 1) under a time-reversible substitution model.

### A2. Factorizing probability into regional contributions

The perturbation theory unfolded in this paper can also be applied, with some extensions, to the entire model defined by the rate operator 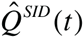 in Eq.(1.1) of Results (*i.e*., Eq.(A1.1)). Because indels are much less frequent than substitutions, a natural way would be to separate 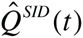 as:

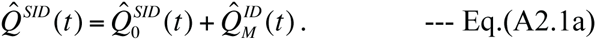

Here

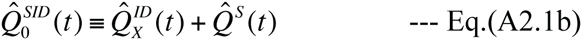

describes the sequence evolution via no indels. And 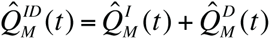, which is the aforementioned extension of Eq.(3.1.1c) of part I (Ezawa, Graur and Landan 2015a), describes the sequence change via an insertion or a deletion. The point is that the *entire* substitution rate operator 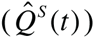 is included in the “unperturbed” part of the rate operator 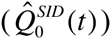, and that *only* insertions/deletions are regarded as “perturbations.” Then, the perturbation expansion of the *entire* stochastic evolution operator, 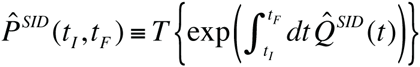, is given by Eq.(3.1.3) of part I with the extension of the state space and with

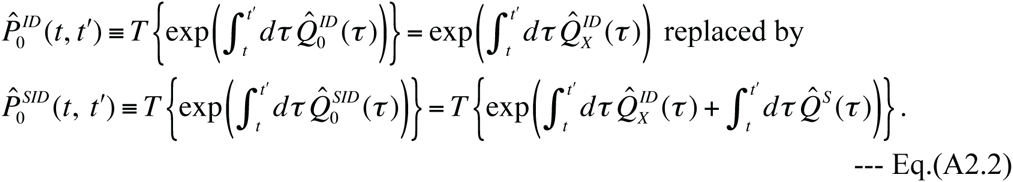

The Eq.(3.1.13) of part I for the probability of a PWA between an ancestral and a descendant sequence, *P*[(*α*(*s^A^*, *s^D^*), [*t*_*I*_, *t*_*F*_])|(*s^A^*, *t*_*I*_)], can also be extended as follows:

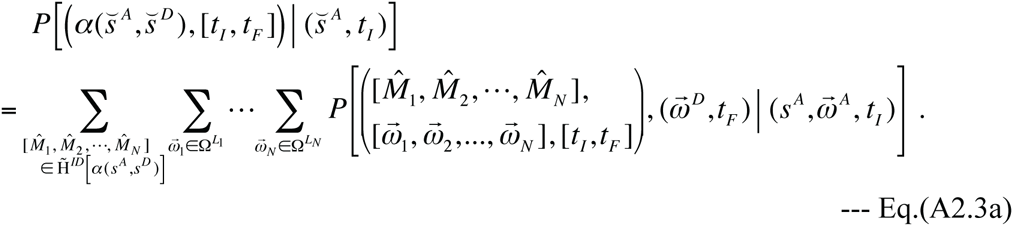

Here we let 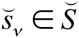 (*v* = 1, 2,…, *N*) be the extended sequence state “at the time of” the event 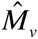, or, more precisely, the state immediately before and after the event if the event is a deletion and an insertion, respectively. We also let 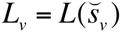 be the number of sites in 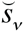. Let 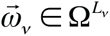 and *s _v_* ∈ *S*, respectively, be the residue state component and the basic component of 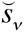. And let 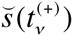 and 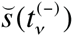, respectively, denote the extended sequence states immediately after and before the time *t*_*v*_ of the event 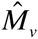 within an evolutionary process. Then, we have 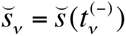 and 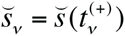 respectively, when 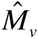 is a deletion and insertion. This choice of 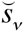 automatically takes account of the summations over 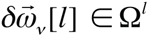 filled in by the accompanying operator 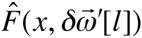 when 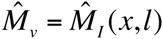. Moreover, we also used the representation 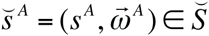, with 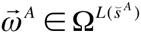, and, similarly, 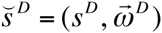. Each summand on the right hand_side of Eq.(A2.3a) represents the probability, conditioned on an ancestral state 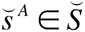 at time *t*_*I*_, that we have the following 3 outcomes at the same time: (1) an indel history 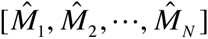 consistent with *α*(*s^A^*, *s^D^*) occurred during the time interval [*t*_*I*_, *t*_*F*_]; (2) an array of residue states 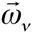 was observed “at the time of” each event 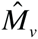; and (3) a descendant array of residue states 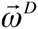 resulted at time *t*_*F*_. The summand is an extension of Eq.(3.1.8b) of part I and is specifically expressed as:

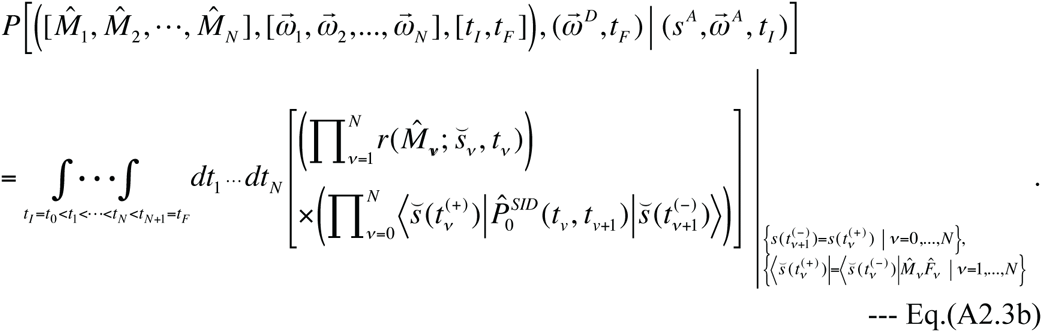

Here 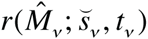 denotes 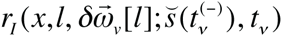 and 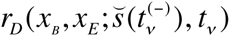 respectively, if 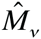 is 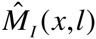 and 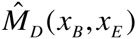. (Here 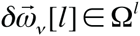 denotes an array of residues filling in the sites inserted by 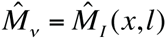. When 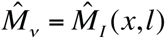, the combination of 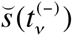, the insertion position (*x*), and 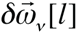 holds information equivalent to that of 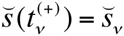.) We also used the notation, 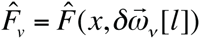 if 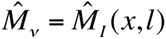, and 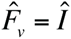 (*i.e*., does nothing) if 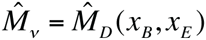. The factor 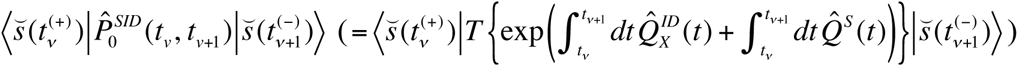 is an extension of 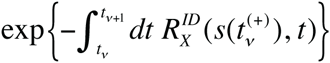 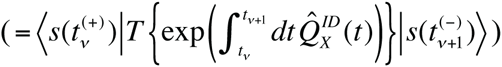 in Eq.(3.1.8b) of part I. (Here, the notation of the latter was changed from the original, to fit in context.) It should be noted that, in the evolutionary process under consideration, the basic state 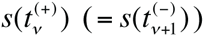 remained unchanged during the open time interval (*t*_*v*_, *t*_v+1_). Thus, during this time interval, the evolutionary changes of the extended state 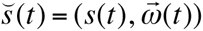 are limited to changes in the residue state 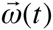 purely via substitutions. In a special case where the indel exit rate, 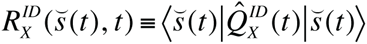, is independent of 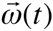 (and thus depends only on *s*(*t*) and *t*), the factor is calculated as:

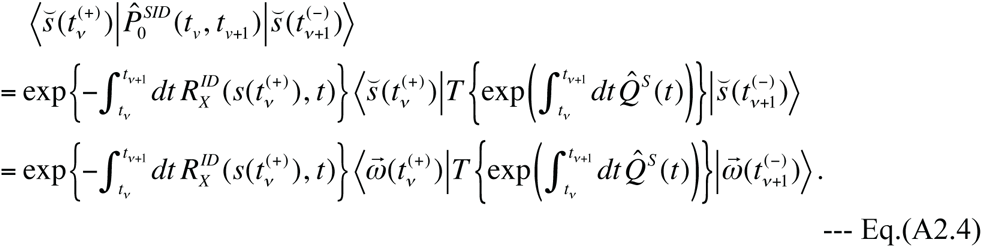

Its only difference from the factor in Eq.(3.1.8b) of part I is the multiplication by the transition probability from 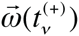 to 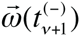 via substitutions. In more general cases where the indel exit rate 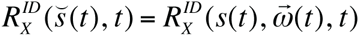 could depend on the residue state 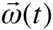, we will decompose the substitution rate operator as 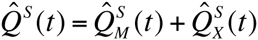 as argued above on the pure substitution case, and follow a line of argument similar to that in Subsection 3.1 of part I. Let 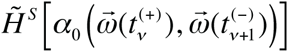 be the set of all substitution histories consistent with the gap-free (*i.e*., trivial) PWA, 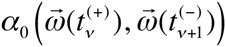, between the residue states at the time boundaries. Then, we get:

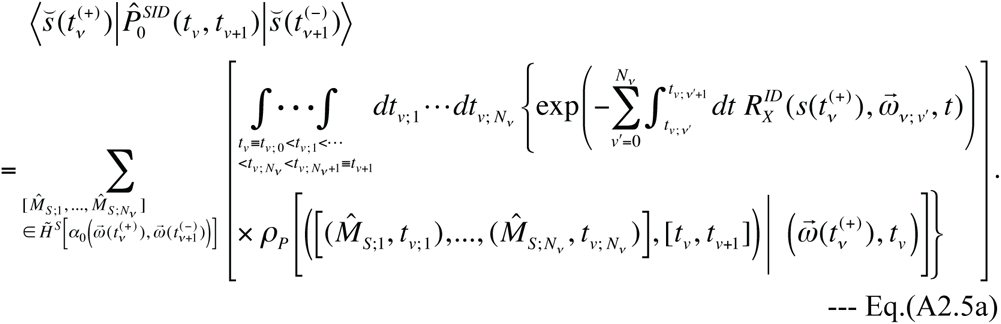

Here, 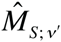(with *v*′ = 1,…, *N*_*v*_) is the *v*′ th substitution event, like 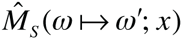 above, in a substitution history during the open time interval (*t*_*v*_, *t*_*v*+1_). *t*_*v*; *v*′_ is the time at which the *v*′ th event occurred. And 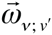 is the residue state of the sequence immediately after the *v*′ th event, with an exception 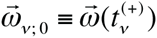. Also, *ρ_P_* […] in Eq.(A2.5a) is the probability density of the substitution process 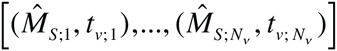, conditioned on the initial residue state 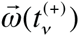. It is formally similar to the integrand in Eq.(3.1.8b) of part I, and is given by:

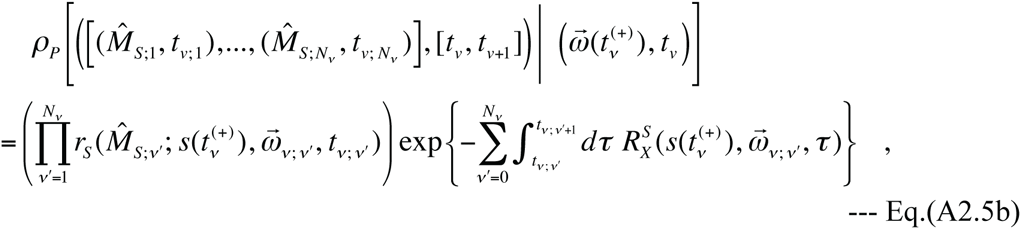

where we used the same notation as in Eq.(A2.5a). The symbol 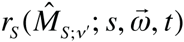 denotes the rate parameter of the substitution 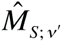 on the extended state 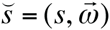 at time *t*. And 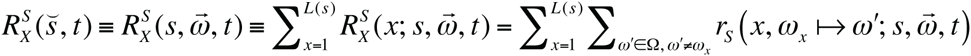 is the substitution exit rate of the extended state 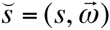 at time *t*. When deriving Eq.(A2.5b), we used the equation: 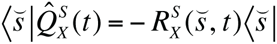. Eq.(A2.5a) suggests that the factor 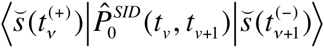 is a weighted summation of 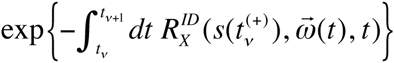 over all substitution processes (each represented by a trajectory 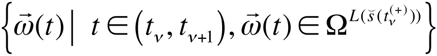 consistent with the (trivial) PWA, 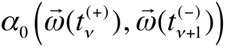, with the weights given by the probability densities of the processes. Because Eq.(A2.5a) supplemented with Eq.(A2.5b) is similar in form to Eq.(3.1.13) of part I supplemented with Eq.(3.1.8b) of part I, a reasoning similar to that in Subsection 4.1 of part I is also applicable when examining the factorability of 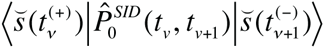. We see that it is factorable into the product of an overall factor and contributions from separate regions if the two conditions are met. (i) The rate parameter 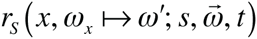 of every substitution 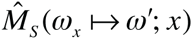 in each region is independent of the portions of the residue states in the other regions. And (ii) the increment of the total exit rate 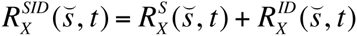, caused by every substitution 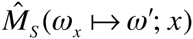 in each region, is independent of the portions of the residue states in the other regions.

Then, substituting the factorized Eq.(A2.5a) for 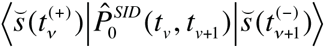 into Eq.(A2.3b), and substituting the result into Eq.(A2.3a), we can examine the factorability of the total probability, 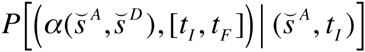, of a PWA between the extended states of the ancestral and descendant sequences. Again, we can follow a line of reasoning similar to that in Subsection 4.1 of part I, with two differences: (a) here, 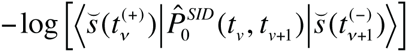 plays the role of 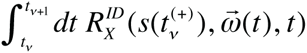 and (b) here, not only the basic states but also the residue states comes into question. Moreover, thanks to the preceding argument, we know that a change in 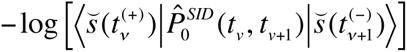 results from a collective effects of the changes in the substitution rates and the total exit rates in Eqs.(A2.5a,b). Thus, we find the following sufficient set of conditions under which 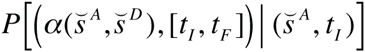 is factorable into the product of an overall factor and contributions from separate regions.

**Condition (I):** In each region, the rate of each of the substitutions and indels is independent of the portions of the extended states in the other regions. And **Condition (II):** In each region, the increment of the total exit rate,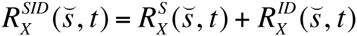, due to each of the substitutions and indels, is independent of the portions of the extended states in the other regions.

As argued in Subsection 3.2 of part I, we could calculate the probability of a given MSA (under a given phylogenetic tree and a given evolutionary model) by assembling the PWAs between the ancestral and descendant sequence states along all branches and by summing over all possible sets of sequence states at internal nodes that are consistent with the MSA. Thus, once we know that the probabilities of PWAs of extended sequence states are factorable, we can also factorize the probability of a given MSA of extended sequence states under a given phylogenetic tree. We can do this by extending the line of arguments unfolded in Subsection 4.2 of part I so that it will incorporate the substitution processes and the resulting residue components of the sequence states. According to such an extended line of arguments, we find that the probability of a given MSA is factorable if the above conditions (I) and (II) holds and, in addition, if we have an extended version of the factorability of the root sequence probability (given in Eq.(4.2.8) of part I):

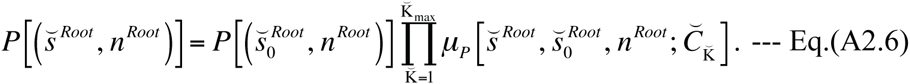

Here 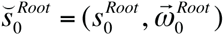 is the extended state of a “reference” root sequence. And the set of regions, 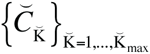, consists not only of the regions potentially accommodating local indel histories ({*C*_K_}_K=1,…,K_max__) but also of the PASs (*i.e*., gapless columns), which can never experience indels but can possibly experience substitutions.

### A3. Factorizing probability into basic and residue components

Thus far, we considered the probability of a given alignment as the summation of the probabilities with consistent indel histories, each of which, in turn, is the summation of the probabilities of substitution histories consistent with the alignment and the indel history (see, *e.g*., Eqs.(A2.3a,b)). The factorization of the probability as discussed above in Subsection A2 could substantially reduce the computational burden of the calculation. However, we will be able to improve the computational efficiency further if we can factorize the entire probability into the product of the “basic” component, which concerns indel processes, and the residue component, which concerns substitution processes (and the initial states of residues). This is because the residue component will then need to be computed only once, instead of as many times as the indel LHSs to be considered. Here we will consider a sufficient set of conditions for such an “indel-substitution factorization.” Because indels are usually at most 1/10 times as frequent as substitutions (*e.g*., Lunter 2007; Cartwright 2009), we will consider, as in Eqs.(A2.3a,b) and Eqs.(A2.5a,b), that each indel history determines the “skeleton” of the alignment that are made up of the basic sequence states at the nodes. Hence we suppose that each substitution history determines the residue states that flesh out the “skeleton” to complete the alignment. First, to simplify the argument, we assume that the probability of the alignment skeleton itself does not depend on the residue states. This assumption is true if the indel rates are independent of the residue states of the sequence immediately before the indels:

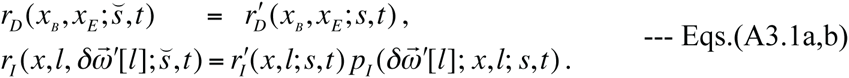

It should be noted that the insertion rates could still depend on the residue states of the *inserted* subsequence through 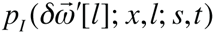, which satisfy 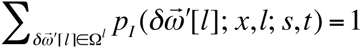. Under the condition Eqs.(A3.1a,b), the indel exit rate 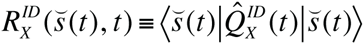 is independent of the residue component of 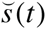, and thus the factor 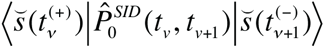 in Eq.(A2.3b) can be factorized as in Eq.(A2.4). Thus, the integrand in Eq.(A2.3b) is reduced to Π*_ID_* × Π*_S_*, with

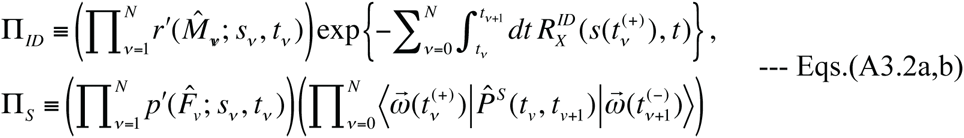

under the same setting and notations as introduced around Eq.(A2.3b). Here 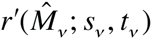 denotes 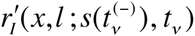 and 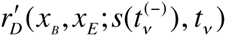, respectively, if 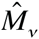 is 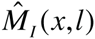 and 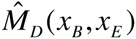. And 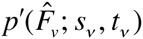 denotes 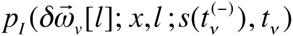 and 1 (unity), respectively, if 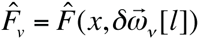 and 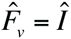. The product, Π*_ID_*, is of the same form as the integrand in Eq.(3.1.8b) of part I (Ezawa, Graur and Landan 2015a), which represents the (residue-independent) probability distribution of an indel process, 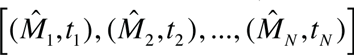. The product, Π*_S_*, is the joint probability of the residue states, 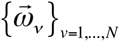, “at the times of” the indel events and of the final residue state 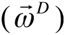, conditioned on the initial residue state 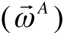 and the indel process. One major difference between Eq.(3.1.13) of part I and its extension, Eq.(A2.3a), is that the latter performs the summation over all possible 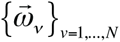. Therefore, if the summation of Π*_S_*’s becomes independent of the indel process, then, it can be factored out of the multiple-time integration in Eq.(A2.3b) and further be factored out of the summation over the consistent indel histories in Eq.(A2.3a). This means that the total probability of a PWA can be factorized into the product of the probability of the “skeleton” of the PWA (as in Eq.(3.1.13) of part I) and the probability of the residue component of the PWA. For the latter, the calculation techniques have been developed well (*e.g*., Felsenstein 1981, 2004; Yang 2006).

Thus the problem is reduced to the condition under which the aforementioned summation of Π_*S*_’s over all 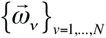 becomes independent of the indel process consistent with the PWA. It would be convenient to categorize the columns in the PWA according to their histories, *i.e*., whether or not they are preserved throughout [*t*_*I*_, *t*_*F*_], and, if not, the time of insertion (*t*_*v*(*I*)_) and/or the time of deletion (*t*_*v*(*D*)_). (If a column has both *t*_*v*(*I*)_ and *t*_*v*(*D*)_, *v* (*D*) < *v* (*D*) should always hold.) And here, we will make a second simplifying assumption, that is, the substitution rates within each block of contiguous columns with a shared history are assumed as independent of the portions of the residue states in the rest of the sequence. Then, the factor 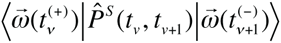 in Eq.(A3.2b) can be factorized into the product of partial probabilities, 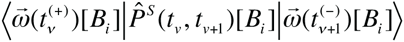, of such blocks with shared histories (denoted as *B*_*i*_ (*i* = 1,…, *I*_*B*_)). And we also assume that 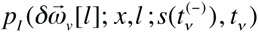 can be factorized into the product of components, denoted as 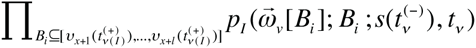. Here each component 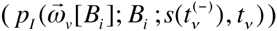 comes from a relevant block, 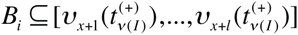. (Note that the blocks are positioned relative to the MSA (or the array of ancestries) but not relative to a particular sequence state.) These two assumptions make the summation of Π*_S_*’s over the intermediate residue states also factorable into the contributions from such blocks. In the following, we consider the contributions from such blocks to the summation of Π*_S_*’s. For this purpose, we broadly classify them into four classes: (1) when *B*_*i*_ was preserved throughout [*t*_*I*_, *t*_*F*_]; (2) when it existed at *t*_*I*_ but was deleted at *t*_*v*(*D*)_; (3) when it was inserted at *t*_*v*(*I*)_ and was deleted at *t*_*v*(*D*)_; when it was inserted at *t*_*v*(*I*)_ and was preserved through *t*_*F*_.

When *B*_*i*_ was preserved throughout [*t*_*I*_, *t*_*F*_], the contribution from the block is:

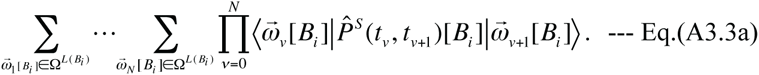

Because 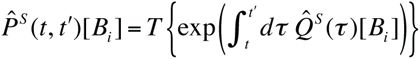 itself is a stochastic evolutionary operator made of the block-wise substitution rate operator 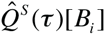, it satisfies the Chapman-Kolmogorov (CK) equation:

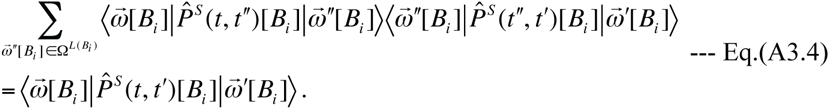

Applying the CK equation successively regarding the intermediate residue states 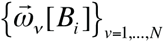, Eq.(A3.3a) is reduced to 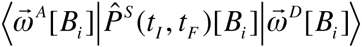, which depends only on the initial and final time points, and the residue states at these time points.

(2) When *B*_*i*_ existed at *t*_*I*_ but was deleted at *t*_*v*(*D*)_, the block’s contribution is:

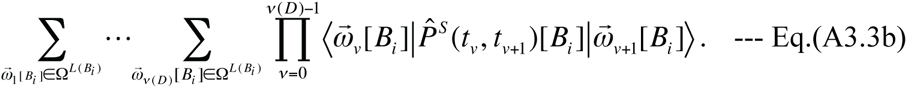

The CK equations apply to the summations over 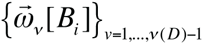, and Eq.(A3.3b) is reduced to 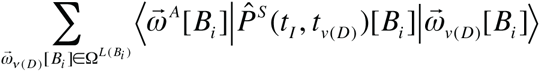. The summand is the probability of 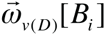 at time *t*_*v*(*D*)_ conditioned on the initial state 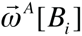. Thus, in this case, the summation Eq.(A3.3b) gives 1 (unity). [*B*_*i*_].

(3) When *B*_*i*_ was inserted at *t*_*v*(*I*)_ and was deleted at *t*_*v*(*D*)_, the block’s contribution is:

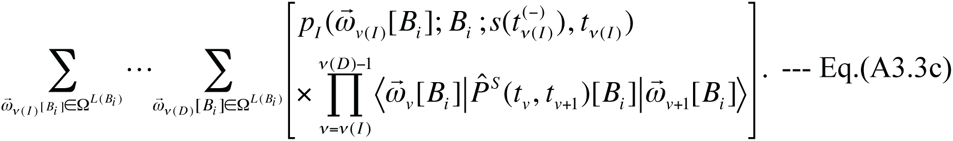

The CK equations apply to the summations over 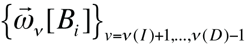, and we have:

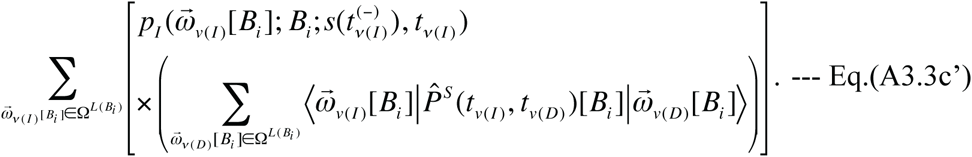

As in case (2), the summation over 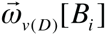’s in the parentheses in Eq.(A3.3c’) gives unity, thus the equation is reduced to 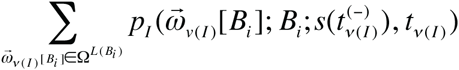. This summation is nothing other than 1 (unity) thanks to the normalization condition of 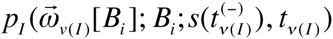. Thus, in conjunction with (2), we see that the contribution from a deleted block to the *conditional* probability of a PWA is always unity, regardless of whether it already existed in the initial sequence or it was inserted.

(4) When *B*_*i*_ was inserted at *t*_*v*(*I*)_ and was preserved through *t*_*F*_, the block’s contribution is:

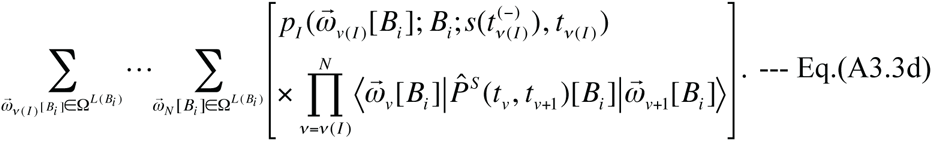

The CK equations apply to the summations over 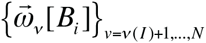, and we have:

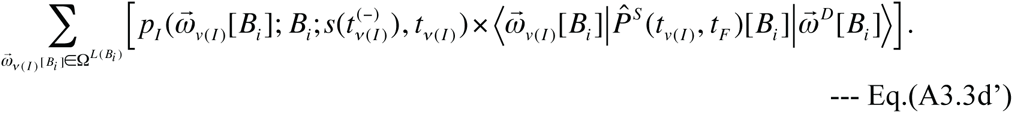

In order for the summation of Π*_S_*’s to be independent of the indel process, Eq.(A3.3d’) needs to be independent of *t*_*v* (*I*)_, especially it needs to be equal to its limit under *t*_*v* (*I*)_ → *t*_*F*_. In this limit, we have 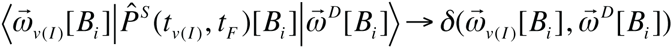, thanks to the defining property of the stochastic evolutionary operator (Eq.(1.1.10b’) of part I). Thus, the aforementioned condition can be expressed as:

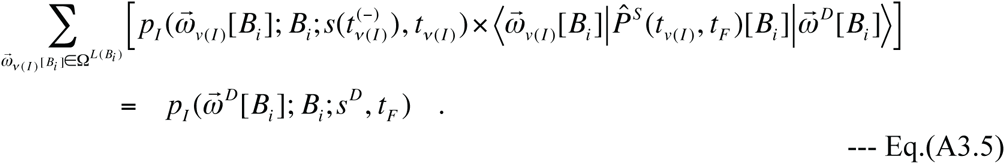

Here we used 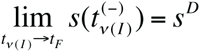. The right hand side of Eq.(A3.5) depends only on the final time, a fixed block, and the portion of the extended sequence state in the block at the final time, and therefore it is totally independent of the indel process, as required.

Thus, let us assume that the four conditions we imposed above are satisfied. Namely, (i) the indel rates are independent of the residue states; (ii) each finite-time evolution probability of the residue state via substitutions is factorable into the product of the probabilities of blocks of particular histories; (iii) the residue state spectrum of each inserted sub-sequence is factorable into the product of block-wise contributions; and (iv) each block-wise spectrum of inserted residue states satisfies Eq.(A3.5). Under these conditions, the summation of Π*_S_*’s (given in Eq.(A3.2b)) over all possible intermediate residue states is expressed as:

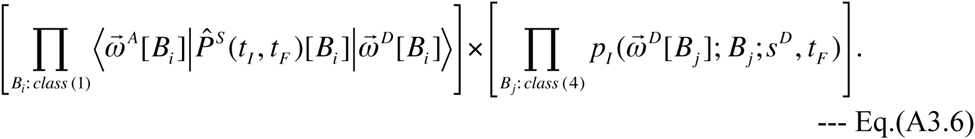

Thus, those sites that didn’t make it through *t*_*F*_ do not contribute to Eq.(A3.6). (More precisely, each of them contributes a trivial multiplication factor of 1 (unity).) Eq.(A3.6) is independent of the indel process that provides the “skeleton,” except the possible dependence on the particular way how the indel process partitions the PWA into blocks. If we consider all indel processes that are consistent with the PWA, there could be a variety of ways of partitioning it into blocks. A simplest way to assure the independence on the way of partitioning the PWA is to assume the following two properties. (1) The stochastic evolutionary operator of substitutions is factorable into the product of site-wise operators: 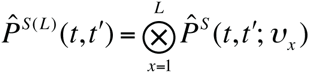. And (2) the residue state spectrum of inserted sub-sequence, 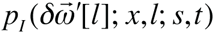 in Eq.(A3.1b), is also factorable into the product of site-wise contributions: 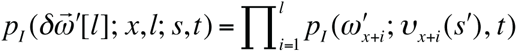. Here, 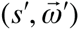 is the extended sequence state immediately *after* the insertion, and we used the ancestries (*v*_*x*_’s) instead of the site numbers (*x*’s) as arguments because the former is invariant through an indel process. Under this assumption, the condition Eq.(A3.5) is reduced to the following single-site condition:

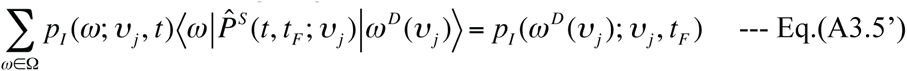

for ^∀^*t* ∈ [*t*_*I*_, *t*_*F*_]. Then, after partitioning the PWA, 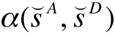, into the columns with ancestries {*v*_1_, *v*_1_,…, *v*_*L* (*α*)}_, where *L*(*α*) is the number of columns in the PWA, Eq.(A3.6) is further reduced to:

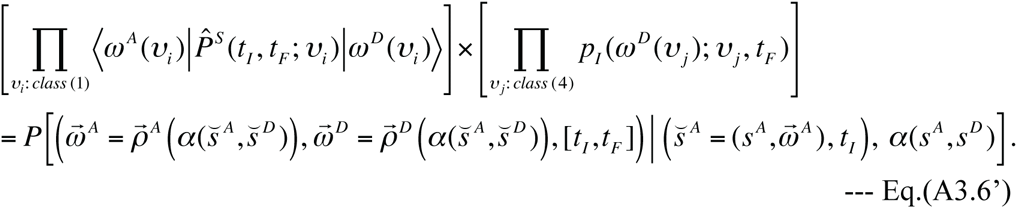

Here *ω^A^*(*v*_*i*_) and *ω*^*D*^(*v*_*i*_), respectively, denote the ancestral and the descendant residues in the PWA column (*i.e*., site) with ancestry *v*, and 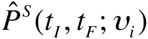 is the single-site evolutionary operator via substitutions in the site with ancestry *v*_*i*_. On the right hand side, 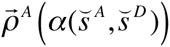 and 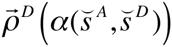, respectively, symbolically represent the vector functions extracting the ancestral and descendant residue states from the PWA of extended sequence states 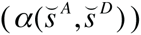. As desired, the left hand side of Eq.(A3.6’) depends only on the ancestral and descendant states, as well as the homology structure of the PWA, and does not depend on any details of the indel processes. Hence, the right hand side follows. Thus, in this case, the summation of Π*_S_*’s can indeed be factored out of the multiple-time integration in Eq.(A2.3b) and also of the summation over all indel histories consistent with the PWA in Eq.(A2.3a). This finally enables us to re-expresses Eq.(A2.3a), supplemented by Eq.(A2.3b), into the form we desire:

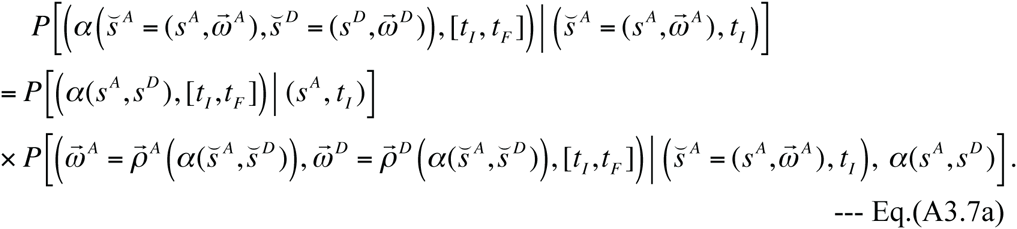

On the right hand side, the second factor is given by Eq.(A3.6’), and the first factor is given by:

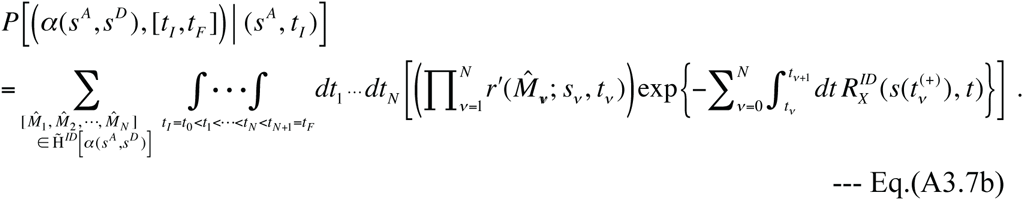

Here we remind the equations, 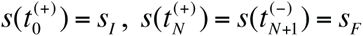, and 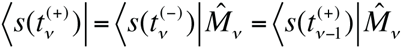 for *v* = 1,…, *N*. Eq.(A3.7b) corresponds exactly to Eq.(3.1.13) of part I supplemented by Eq.(3.1.8b) of part I, which gives the probability of the “skeleton” *α*(*s^A^*, *s^D^*) of the PWA 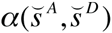, conditioned on the basic ancestral state *s^A^* at the initial time, due to indel processes.

The above line of arguments implies that it would be very difficult, even if it is possible at all, to factorize a whole probability of a PWA into the product, Eq.(A3.7a), of the basic component (Eq.(A3.7b)) and the residue component (Eq.(A3.6’)), *unless* the residue component of the stochastic evolution operator is factorable into the product of column-wise operators as in Eq.(A1.3), or *unless* the indel rate parameters can be expressed as in Eqs.(A3.1a,b), where 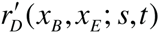 and 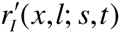 are independent of the residue states. Nevertheless, even if either of these conditions is violated, the whole probability of a PWA is still factorable into the product of the contributions from some separated regions, if the rate parameters of indels and substitutions in each region are independent of the portions of the extended sequence state outside of the region, as argued in Subsection A2. Therefore, even if the residue states at some sites have substantial impacts on the indel rates and/or the substitution rates, if such effects are localized in some narrow regions, we could first factorize the entire PWA probability into the product of regional contributions and then factorize most of such regional contributions into the basic and the substitution components. Then, we could separately handle the small portions whose “indel-substitution factorizations” are not possible. This way, the computational burden may still be mitigated considerably. Such a situation might apply to some mutagenic and/or functional motifs that are scattered along the sequence and which show quite rapid turnover. (If a motif is strongly conserved, in contrast, its effect will be well approximated by the *ancestry* dependence, instead of the residue dependence, of the rate parameters, and thus the “indel-substitution factorization” holds at least approximately.)

Now, let us assume the aforementioned conditions for the indel-substitution factorization of the conditional probability of PWAs, *i.e*., that the indel rate parameters are of the forms in Eqs.(A3.1a,b) and that the substitution evolutionary operator is factorable into the product of the column-wise operators as in Eq.(A1.3). Under such conditions, we will examine whether the probability of a given MSA is also factorable into the indel (*i.e*., “basic”) and substitution (*i.e*., residue) components. In this case, we generalize Eqs.(3.2.13a,b’) of part I to the probability of an “extended *α* MSA,” *i.e*., an alignment of multiple extended sequence states, 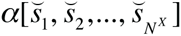. The set of all sets of basic states at internal nodes consistent with the MSA, *i.e*., 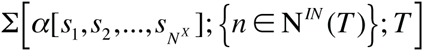 in Eq.(3.2.13a) of part I, can be extended by filling in each of the sets of internal basic states with all possible residue states. Thus, the extended set is expressed as:

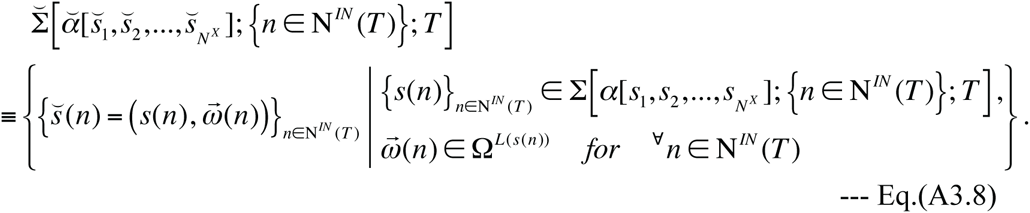

Hence, we have the extended version of Eq.(3.2.13a) of part I as follows:

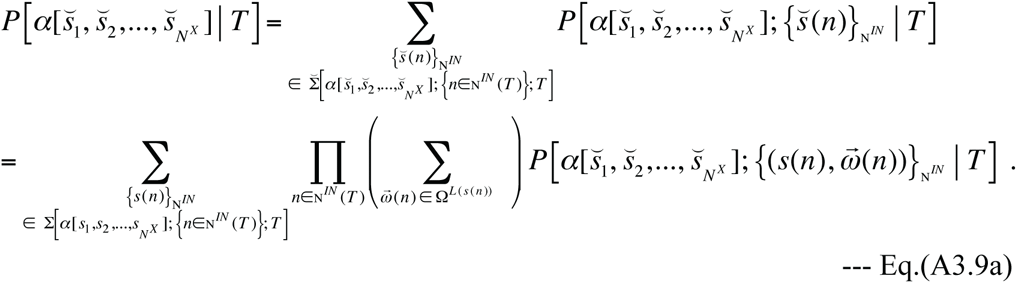

Here,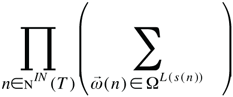 represents the multiple summations over all possible sets of residue states at internal nodes. More precisely, in each possible set, 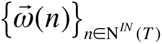, each component state 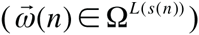 fills in a fixed basic state (*s*(*n*)) at each internal node (*n* ∈ N*^IN^*(*T*)). And the probability, 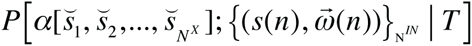, is given by an extended version of Eq.(3.2.13b’) of part I:

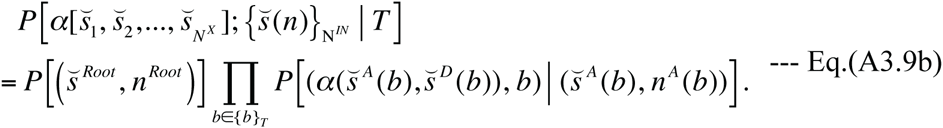

Here, as below Eq.(3.2.13b’) of part I, 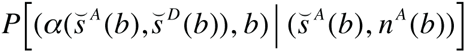 denotes the probability of a PWA between the *extended* states at the ancestral and descendant nodes of branch *b*. Under the present assumptions, each of such probabilities can be factorized into the basic and residue components, as in Eq.(A3.7a). In addition, we assume that the root state probability is also factorable as:

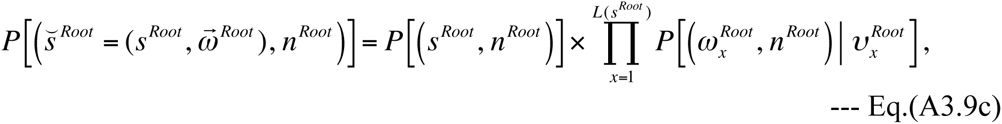

where 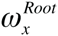 and 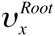 denote the residue and basic components, respectively, at the *x* th site of the root sequence. By substituting Eqs.(A3.7a, 9c) into Eq.(A3.9b), we see that the summand in Eq.(A3.9b), *i.e*., 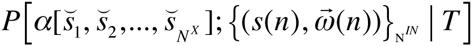 is factorized as:

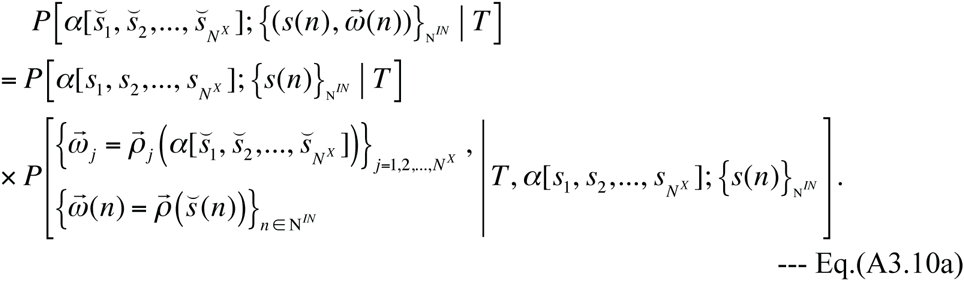

Here, 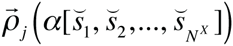 (with *j* = 1, 2,…, *N*^*X*^) symbolically represents the vector function extracting the residue state of 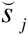 in the MSA of extended sequence states 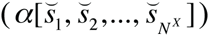. And 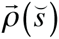 represents the vector function extracting the residue state of an extended sequence state 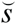. On the right hand side of Eq.(A3.10a), the first factor, *P*[*α*[*s*_1_, *s*_2_,…, *s_N^X^_*]; {*s*(*n*)}_N^*IN*^_| *T*], is given by an equation that exactly corresponds to Eq.(3.2.13b’) of part I. Meanwhile, the second factor is given by:

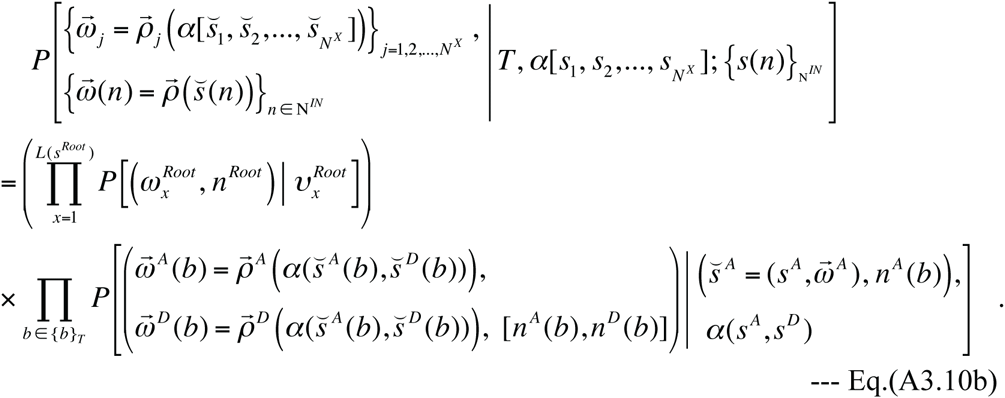

Here each conditional probability in Π_*b*∈{*b*}_*T*__ (…) is given by the left hand side of Eq.(A3.6’) slightly modified to fit the evolution along each branch *b*. Thus, the right hand side of Eq.(A3.10b) can be re-expressed as a product over contributions from single columns of 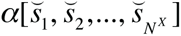, where each single column’s contribution is also a product of terms coming from different branches and nodes. If we can show that the summation of the right hand side of Eq.(A3.10b) over all possible internal residue states is independent of particular details of the basic states at internal nodes as long as they are consistent with the MSA, then, the summation can be factored out of the summation over Σ[*α*[*s*_1_, *s*_2_,… *s_N^X^_*]; {*n* ∈ *N^IN^* (*T*)}; *T*] on the right hand side of Eq.(A3.9a). If so, the total probability of a given extended MSA, 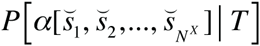 given by Eq.(A3.9a), can be factorized into the basic and residue components:

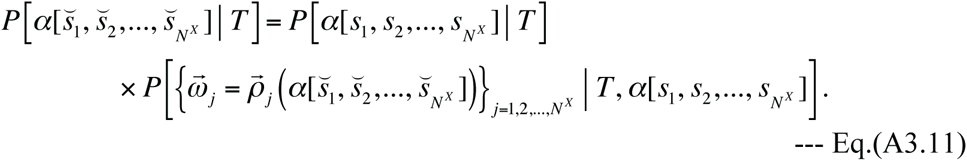

Here *P*[*α* [*s*_1_, *s*_2_,…, *s_N^X^_* | *T*] is given by an equation exactly corresponding to Eq.(3.2.13a) of part I. And 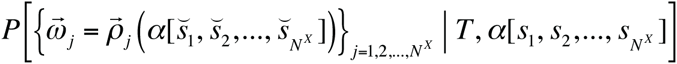 is the summation of Eq.(A3.10b) over all possible internal residue states filling in a fixed set of basic internal states ({*s*(*n*)}_N^*IN*^_) consistent with the MSA. In the following, we will show that this summation is indeed independent of the details of {*s*(*n*)}_N^*IN*^_.

First, because Eq.(A3.10b) with the reverse-substitution by Eq.(A3.6’) is a product of single-site contributions, we can sort it into a product of column-wise probabilities (*P*(*v*_*i*_)’s) over all MSA columns, which are assigned ancestries *v*_1_, *v*_2_,…, *v*_*L* (*α*)_ (*L*(*α*) is the number of columns in the MSA):

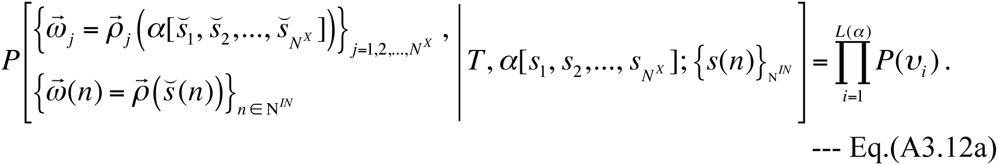

Because each MSA column, *i.e*., each site with a given ancestry (*v*_*i*_), can experience *at most* one insertion, each *P*(*v*_*i*_) can be broadly classified into two forms, as follows.

(1) When the site did not experience an insertion, the site already existed at the root node. Thus, considering Eq.(A3.6’) and Eq.(A3.9c), we have:

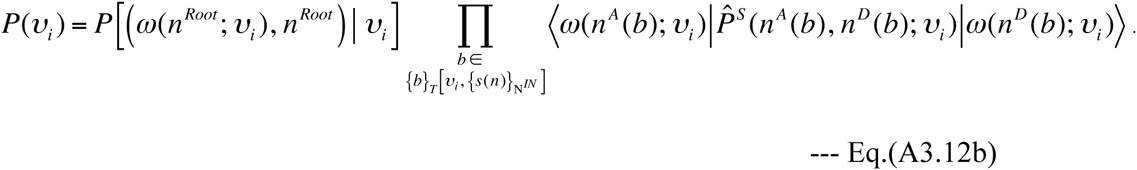

Here, *ω*(*n*; *v*_*i*_) denotes the residue state at the site with the ancestry *v*_*i*_ in the extended sequence state at the node *n*. And {*b*}*_T_* [*v*_*i*_; {*s*(*n*)}_N^*IN*^_] is the set of branches through which the site *v*_*i*_ continued to exist (*i.e*., the site experienced no indel event along the branch), given a set of the internal node states ({*s*(*n*)}_N^*IN*^_). (2) When the site experienced an insertion along a branch *b*_*I*_, we have:

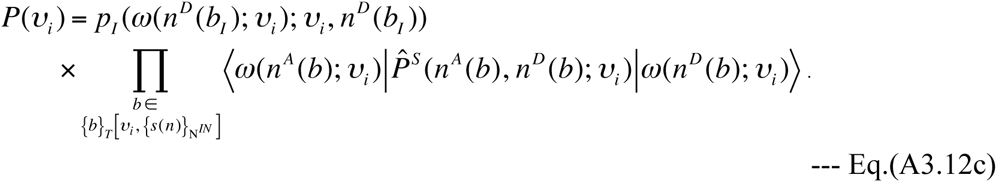

Here, inevitably, {*b*}*_T_* [*v*_*i*_; {*s*(*n*)}_N^*IN*^_] consists solely of some, but not necessarily all, of the descendant branches of *b*_*I*_.

Now, in a MSA column (with ancestry *v*_*i*_), consider the summation of *P*(*v*_*i*_)’s over all possible residue states at relevant internal nodes. Let N*^IN^*[*v*_*i*_; {*s*(*n*)}_N^*IN*^_] be the set of such internal nodes. According to the phylogenetic correctness condition (*e.g*., Childelevitch et al. 2006) (as mentioned near the bottom of Subsection 3.2 of part I), the union of N*^IN^*[*v*_*i*_; {*s*(*n*)}_N^*IN*^_], the set of external nodes with the site of ancestry *v*_*i*_ (N*^X^*[*v*_*i*_]), and {*b*}_T_[*v*_*i*_; {*s*(*n*)}_N^*IN*^_] always forms a single, connected web in the tree (see the lower part of Subsection 3.4 of part I, and Figures 4 and 7 of part I). The minimum web among such webs is given by the Dollo parsimonious indel history (Farris 1977), and is a union of N*^X^*[*v*_*i*_] and the shortest paths, each of which connects a pair of nodes in N*^X^*[*v*_*i*_]. Other webs consistent with the MSA column are formed by continuously extending one or more paths from the minimum web. There are two types of web-extension: upward (toward the root) and downward (toward, but always short of, the external nodes not in N*^X^*[*v*_*i*_]). Downward extensions could branch off, as long as the branches do not reach any external nodes. We first handle downward extensions and then handle upward extensions. At each lower-tip of a downward extension, we always encounter a summation of single conditional probabilities, such as 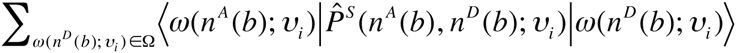, which always gives 1 (unity). After repeating this type of summations, the residue state probabilities at nodes on each downward extension (except its origin belonging to the minimum web) leave no effects on the column-wise MSA probability via substitutions. Next, at each upper-tip of an upward extension, we encounter the following summation: either 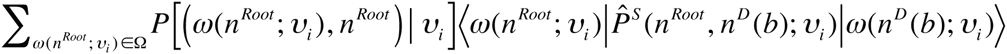 if the tip is the root node (*i.e.*, *n^A^* (*b*) = *n^Root^*), or 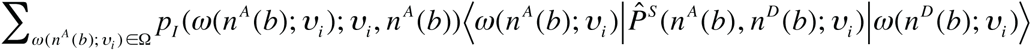 otherwise (*i.e*., if *n^A^*(*b*) = *n^D^*(*b*)). Each of them is a summation over the initial states, and each summand is the product of an “initial probability” and a single probability conditioned on the initial state. Thanks to Eq.(A3.5’), the latter type of summation can be performed, yielding *p*_I_ (*ω*(*n^D^*(*b*); *v*); *v*, *n^D^*(*b*)). The former type of summation can also be performed *if we additionally assume* the following equation:

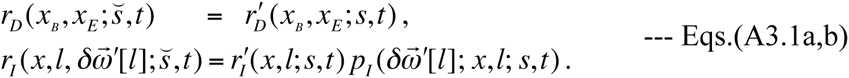

Thus, if Eq.(A3.13) holds, every upward extension can be receded down to its origin (*n^Ori^*) belonging to the minimum web, providing *p*_*I*_ (*ω*(*n^Ori^*; *v*_*i*_); *v*_*i*_, *n^Ori^*). Finally, consider the contribution from a “null site” that is not kept at any external nodes. In this case, after successively performing the summations at the lower-tips of the downward extensions, we are always left with either Σ_*ω*′∈Ω_ *p* [(*ω′*, *n^Root^*)| *v*_*i*_] or Σ_*ω*′∈Ω_ *p*_*I*_ (*ω*′; *v*_*i*_, *n*). Each of them always yields 1 (unity). Putting together all these arguments, we see that the probability of the residue states of a MSA column under any indel history can be reduced to that under the Dollo parsimonious indel history. This means that the residue component of the MSA probability, which is the summation of Eq.(A3.10b) over all possible residue states at internal nodes, is indeed independent of the basic sequence states at internal nodes, {*s*(*n*)}_N^*IN*^_. Thus, under the assumptions of Eqs.(A3.1a,b), the column-wise factorization of the substitution evolutionary operators, Eq.(A3.5’) and Eq.(A3.13), the MSA probability can indeed be factorized into the basic and residue components, as in Eq.(A3.11).

### A4. Pursuing further biological realism

Eq.(A3.5’) and Eq.(A3.13) in Subsection A3 are *non-equilibrium* generalizations of the famous detailed-balance condition, 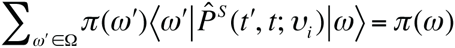, for a time-reversible substitution model with the equilibrium residue frequencies {*π* (*ω*)}_*ω*∈Ω_ and the assumption that the residue content of the inserted subsequences is also given by {*π* (*ω*)}_*ω* ∈Ω_. These widely accepted assumptions played important roles to facilitate the calculations in the past studies with evolutionary models incorporating both substitutions and indels (*e.g*., Thorne et al. 1991, 1992; Miklós et al. 2004; Rivas and Eddy 2008). However, even if generalized to Eq.(A3.5’) and Eq.(A3.13), they may still be too restrictive to accommodate some biologically realistic features. For example, when transposable elements (*e.g*., Morgante et al. 2007; Chalopin et al. 2015) or foreign DNA sequences (*e.g*., Waterhouse and Russell 2006) are inserted, the residue content of such inserted sequences is likely to be substantially different from the residue content of the genome that underwent the insertions. It remains to be seen whether we can further relax the conditions to accommodate such situations while keeping the “indel-substitution factorization” enabled. Recently, Lèbre and Michel (2010, 2013) developed some analytical models to examine the effects of the base composition of inserted sequences on the evolution of the base composition of an entire genome or of its subset. It might be interesting to see if their methods are applicable to the issue at hand.

Another potentially important biologically realistic feature is the observation by some studies that the substitution rate increases at sites surrounding insertions/deletions (*e.g.*, Tian et al. 2008; De and Babu 2010). If the incremental substitutions occurred simultaneously or almost simultaneously with the indel events, this feature could be formally incorporated into our theoretical framework by “dressing” each indel operator term in the action of the rate operator,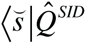, with substitution operators. The deletion operator, 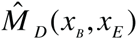, could be replaced with:

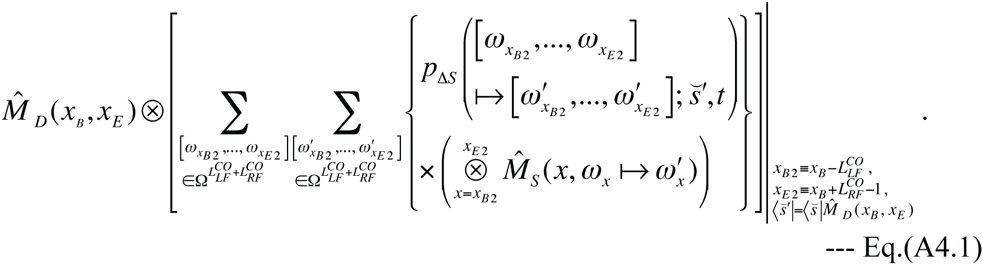

Here 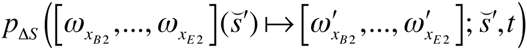 is the probability that the residue states of the intermediate state, 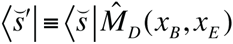, was replaced as indicated, and satisfies 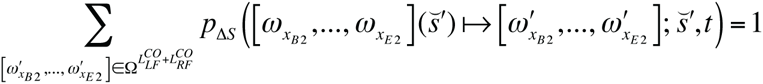. For notational convenience, we have also set 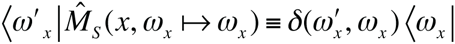, where the subscript *x* in *ω_x_* and *ω*′ *_x_* indicates that they are residue states at the *x* th site. The 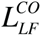 and 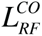 are the “cut-off” lengths of the left-flanking and right-flanking regions, respectively, that could accommodate the incremental substitutions. The cut-offs were introduced just for convenience. If we can assume that the incremental substitution at each site is independent of those at the other sites, Eq.(A4.1) is reduced to:

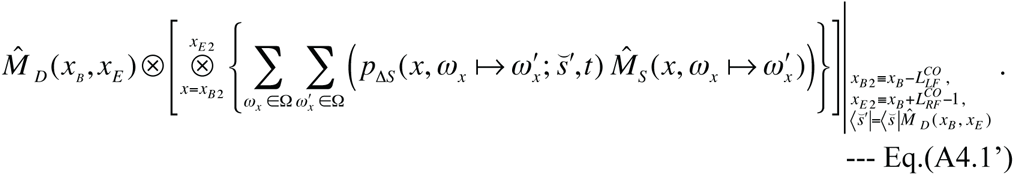

Here the site-wise incremental substitution probability, 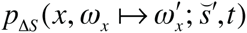, satisfies the equation, 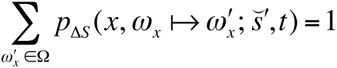. We can also “dress” the insertion operator, 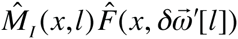, in a similar manner. The expression of a dressed insertion operator becomes bulkier than Eq.(A4.1), and thus is omitted here. Once the incremental substitutions are introduced as in Eq.(A4.1), we cannot easily perform the “indel-substitution factorization” of the alignment probabilities, because the expected number of substitutions increases with the number of indels in a local history.

Therefore, if the incremental substitutions occur commonly along the genome, the above line of arguments is no longer applicable for the separation of basic and residue components of the *entire* alignment probability. Nevertheless, the alignment probability may still be factorable into the product of local contributions, possibly with some modifications in the arguments (in Section 4 of part I (Ezawa, Graur and Landan 2015a)) and the models (given in Section 5 of part I). It remains to be seen whether we can still factorize the alignment probability into the basic and residue components by substantially extending and/or modifying the arguments given in this subsection. Unless we can, one solution might be to develop an “effective substitution model” that takes beforehand account of the effects of such incremental substitutions in the vicinity of indels (including invisible ones).

